# Frequent non-allelic gene conversion on the human lineage and its effect on the divergence of gene duplicates

**DOI:** 10.1101/135152

**Authors:** Arbel Harpak, Xun Lan, Ziyue Gao, Jonathan K. Pritchard

## Abstract

Gene conversion is the copying of genetic sequence from a “donor” region to an “acceptor”. In non-allelic gene conversion (NAGC), the donor and the acceptor are at distinct genetic loci. Despite the role NAGC plays in various genetic diseases and the concerted evolution of gene families, the parameters that govern NAGC are not well-characterized. Here, we survey duplicate gene families and identify converted tracts in 46% of them. These conversions reflect a large GC-bias of NAGC. We develop a sequence evolution model that leverages substantially more information in duplicate sequences than used by previous methods and use it to estimate the parameters that govern NAGC in humans: a mean converted tract length of 250bp and a probability of 2.5×10^−7^ per generation for a nucleotide to be converted (an order of magnitude higher than the point mutation rate). Despite this high baseline rate, we show that NAGC slows down as duplicate sequences diverge—until an eventual “escape” of the sequences from its influence. As a result, NAGC has a small average effect on the sequence divergence of duplicates. This work improves our understanding of the NAGC mechanism and the role that it plays in the evolution of gene duplicates.

## Background

As a result of recombination, distinct alleles that originate from the two homologous chromosomes may end up on the two strands of the same chromosome. This mismatch (“heteroduplex”) is then repaired by synthesizing a DNA segment to overwrite the sequence on one strand, using the other strand as a template. This process is called gene conversion.

Although gene conversion is not an error but rather a natural part of recombination, it can result in the non-reciprocal transfer of alleles from one sequence to another, and can therefore be thought of as a “copy and paste” mutation. Gene conversion typically occurs between allelic regions (allelic gene conversion, AGC) [46]. However, *non-allelic* gene conversion (NAGC) between distinct genetic loci can also occur when paralogous sequences are accidently aligned during recombination because they are highly similar [11]—as is often the case with young tandem gene duplicates [27].

NAGC is implicated as a driver of over twenty diseases [6, 11, 10]. The transfer of alleles between tandemly duplicated genes—or pseudogenes—can cause nonsynonymous mutations [22, 67], frameshifting [51] or aberrant splicing [40]—resulting in functional impairment of the acceptor gene. A recent study showed that alleles introduced by NAGC are found in 1% of genes associated with inherited diseases [10].

NAGC is also considered to be a dominant force restricting the evolution of gene duplicates [48, 16, 21]. It was noticed half a century ago that duplicated genes can be highly similar within one species, even when they differ greatly from their orthologs in other species [58, 57, 8, 38]. This phenomenon has been termed “concerted evolution” [72]. NAGC is an immediate suspect for driving concerted evolution, because it homogenizes paralogous sequences by reversing differences that accumulate through other mutational mechanisms [58, 57, 48, 50]. Another possible driver of concerted evolution is natural selection. Both purifying and positive selection may restrict sequence evolution to be similar in paralogs [26, 64, 59, 27, 16, 60, 41, 20]. Importantly, if NAGC is indeed slowing down sequence divergence, it puts in question the fidelity of molecular clocks for gene duplicates [27, 9]. In order to develop expectations for sequence and function evolution in duplicates, we must characterize NAGC and its interplay with other mutations.

In attempting to link NAGC mutations to sequence evolution, we need to know two key parameters: (i) the rate of NAGC and (ii) the converted tract length; These parameters have been mostly probed in non-human organisms with mutation accumulation experiments limited to single genes—typically artificially inserted DNA sequences [33, 43]. The mean tract length has been estimated fairly consistently across organisms and experiments to be a few hundred base pairs [42]. However, estimates of the rate of NAGC vary by as much as eight orders of magnitude [71, 69, 61, 33, 39]—presumably due to key determinants of the rate that vary across experiments, such as genomic location, sequence similarity of the duplicate sequences and the distance between them, and experimental variability [55, 43]. Alternatively, evolutionary-based approaches[26, 53] tend to be less variable: NAGC has been estimated to be 10-100 times faster than point mutation in *Saccharomyces cerevisiae* [62], *Drosophila melanogaster* [65, 2] and human [26, 52, 7, 25]. These estimates are typically based on single loci (but see [14, 31]). Recent family studies [70, 19, 47] have estimated the rate of AGC to be 5.9 × 10^−6^ per bp per generation. This is likely an upper bound on the rate of NAGC, since NAGC requires a misalignment of homologous chromosomes during recombination, while AGC does not.

Here, we estimate the parameters governing NAGC with a novel sequence evolution model. Our method is not based on direct empirical observations, but it leverages substantially more information than previous experimental and computational methods: we use data from a large set of segmental duplicates in multiple species, and exploit information from a long evolutionary history. We estimate that the rate of NAGC in newborn duplicates is an order of magnitude higher than the point mutation rate in humans. Surprisingly, we show that this high rate does not necessarily imply that NAGC distorts the molecular clock.

## Results

To investigate NAGC in duplicate sequences across primates, we used a set of gene duplicate pairs in humans that we had assembled previously [36]. We focused on young pairs where we estimate that the duplication occurred after the human-mouse split, and identified their orthologs in the reference genomes of chimpanzee, gorilla, orangutan, macaque and mouse. We required that each gene pair have both orthologs in at least one non-human primate and exactly one ortholog in mouse. Since our inference methods implicitly assume neutral sequence evolution, we focused our analysis on intronic sequence at least 50bp away from intron-exon junctions. After applying these filters, our data consisted of 97,055bp of sequence in 169 intronic regions from 75 gene families (**SI Appendix**).

We examined divergence patterns (the partition of alleles in gene copies across primates) in these gene families. We noticed that some divergence patterns are rare and clustered in specific regions. We hypothesized that NAGC might be driving this clustering. To illustrate this, consider a family of two duplicates in human and macaque which resulted from a duplication followed by a speciation event—as illustrated in **Fig. 1B** (“Null tree”). Under this genealogy, we expect certain divergence patterns across the four genes to occur more frequently than others. For example, the grey sites in **Fig. 1C** can be parsimoniously explained by one substitution under the null genealogy. They should therefore be much more common than purple sites, as purple sites require at least two mutations. However, if we consider sites in which a NAGC event occurred after speciation (**Fig. 1A** and “NAGC tree” in **Fig. 1B**), our expectation for variation patterns changes: now, purple sites are much more likely than grey sites.

**Figure 1:**
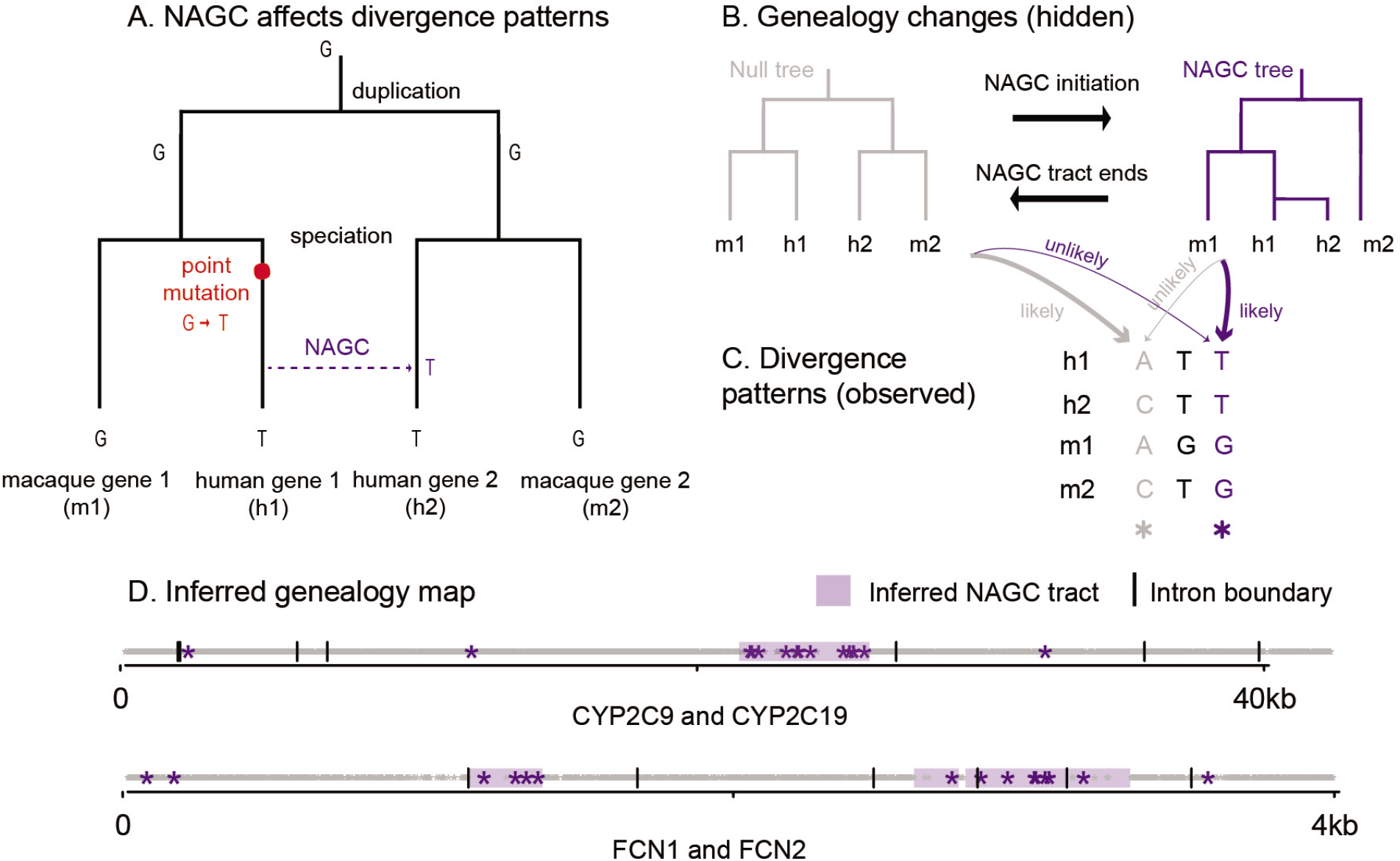
Non-allelic gene conversion (NAGC) alters divergence patterns. **(A)** NAGC can drive otherwise rare divergence patterns, like the sharing of alleles between paralogs but not orthologs. **(B)** An example of a local change in genealogy, caused by NAGC. **(C)** examples of divergence patterns in a small multigene family. Some divergence patterns—such as the one highlighted in purple—were both rare and spatially clustered. We hypothesized that underlying these changes are local changes in genealogy, caused by NAGC. **(D)** Genealogy map (null genealogy marked by white, NAGC by purple tracts) inferred by our Hidden Markov Model (HMM) based on observed divergence patterns (stars). Two different gene families are shown. For simplicity, only the most informative patterns (purple and grey sites, as exemplified in panel C) are plotted.

### Mapping recent NAGC events

We developed a Hidden Markov Model which exploits the fact that observed local changes in divergence patterns may point to hidden local changes in the genealogy of a gene family (**Fig. 1B,C**). In our model, genealogy switches occur along the sequence at some rate; the likelihood of a given divergence pattern at a site then depends only on its own genealogy and nucleotide substitution rates. Our method is similar to others that are based on incongruency of inferred genealogies along a sequence [4, 29, 68], but it is model-based and robust to substitution rate variation across genes (**SI Appendix**).

We applied the HMM to a subset of the gene families that we described above: families of four genes consisting of two duplicates in human and a non-human primate. Since the HMM assumes that the duplication preceded the speciation, we required that the overall intronic divergence patterns support this genealogy, using the software *MrBayes* [24]. This requirement decreased the number of gene families considered to 39.

Applying our HMM, we identified putatively converted tracts in 18/39 (46%) of the gene families, affecting 25.8% of intronic sequence (**Fig. 2A**; see complete list of identified tracts in **Files S1-4**). Previous studies estimate that only several percent of the sequence is affected by NAGC, but the definition of “affected sequence” statistic is arguably method-dependendent and therefore not directly comparable [28, 14, 12]. **Fig. 1D** shows an example of the maximum likelihood genealogy maps for two gene families. The average length of the detected converted tracts is 880bp (**Fig. 2B**). As previously discussed for other methods [43], this is likely an overestimate of the mean tract length of NAGC events, because some identified NAGC tracts result from multiple NAGC events occurring in close proximity (**SI Appendix; Fig. S2**).

**Figure 2:**
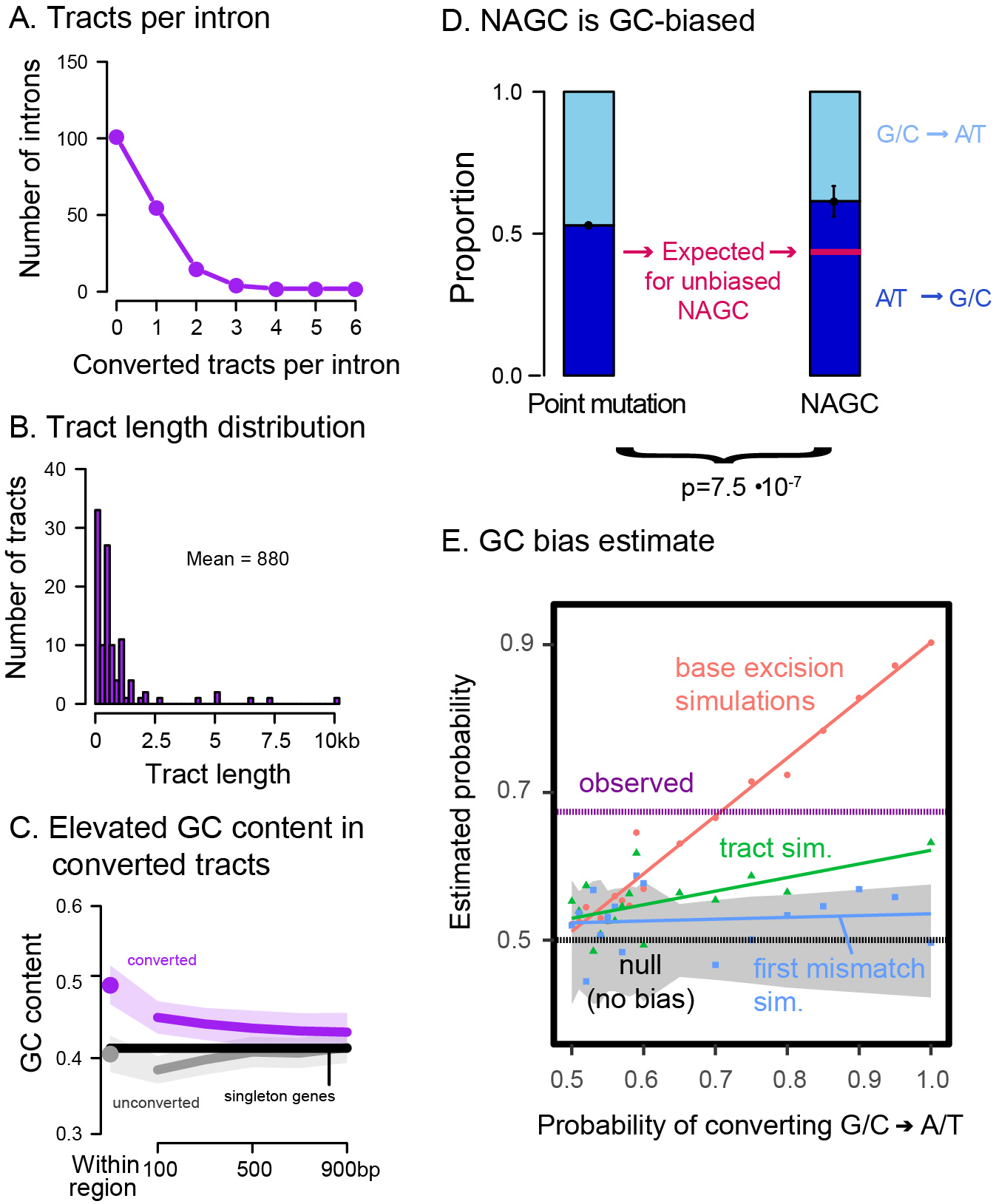
Properties of HMM-inferred converted tracts. **(C)** The purple dot shows the average GC content in converted regions. The grey dot shows the average for random unconverted regions, matched in length and within the same gene as the converted regions. The lines show GC content for symmetric 200bp bins centered at the respective regions (excluding the focal tract). Shaded regions show 95% confidence intervals. Black line shows the intronic average for human genes with no identified paralogs. **(D)** In purple sites (**Fig. 1C**) that are most likely to be a direct result of NAGC (right bar), 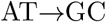 substitutions are significantly more common than 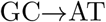 substitutions. The left bar shows the estimated proportion of 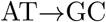 substitutions through point mutations and AGC in unconverted regions, which we used to derive the expected proportion for unbiased NAGC (pink line) after accounting for their different GC contents. Black widgets show two standard errors around the point estimates. **(E)** Point estimate of GC bias. The purple dotted line shows the estimated probability of resolving a GC/AT heteroduplex in favor of the G/C allele. The color dots show simulation results under three different mechanistic models of biased gene conversion. Color lines show linear fits. The grey-shaded area is a 95% binomial confidence interval for the “tract” model with no GC bias.

When an AT/GC heteroduplex DNA arises during AGC, it is preferentially repaired towards GC alleles [15, 49]. We sought to examine whether the same bias can be observed for NAGC [18, 15, 3, 44]. We found that converted regions have a high GC content: 48.9%, compared with 39.6% in matched unconverted regions (*p* = 4 × 10^−5^, two-sided t-test; **Fig. 2C**). This base composition difference has been previously observed for histone paralogs [18]. However, the difference could either be a driver and/or a result of NAGC. To test whether NAGC preferentially repairs AT/GC heteroduplexes towards GC, we focused on sites that carry the strongest evidence of nucleotide substitution by NAGC—these are the sites with the “purple” divergence pattern as before (**Fig. 1C**). Using a parsimony consideration, we inferred the directionality of such substitutions involving both weak (A/T) and strong (G/C) nucleotides. We found that 61% of these changes were weak to strong changes, compared with an expectation of 44% through point mutation differences and GC-biased AGC alone (exact binomial test *p* = 7.5 × 10^−7^ and see **SI Appendix**; **Fig. 2D**). We estimate that this observed difference corresponds to a probability of 67.3% in favor of strong alleles when correcting strong/weak heteroduplexes. Our estimate agrees with the GC bias estimated for AGC [70, 19]. Among several possible repair mechanisms that could underly biased gene conversion that we consider in a simulation study [37, 1], the most likely to underlie such a large bias is the base excision repair mechanism—in which the choice of strand to repair is independent for each heteroduplex (**SI Appendix; Fig. 2E**).

The power of our HMM is likely limited to recent conversions, where local divergence patterns show clear disagreement with the global intron-wide patterns; it is therefore applicable only in cases where NAGC is not so pervasive that it would have a global effect on divergence patterns [42, 5]. Next, we describe a method that allowed us to estimate NAGC parameters without making this implicit assumption.

### NAGC is an order of magnitude faster than point mutation

To estimate the rate and the tract length distribution of NAGC, we developed a two-site model of sequence evolution with point mutation and NAGC (Methods). This model is inspired by the rationale that guided Hudson [23] and McVean et al. [45] in estimating recombination rates: while computing the full likelihood of a sequence evolving through both point mutation and NAGC is intractable, we were able to model the likelihood of the observed divergence between paralogs at a pair of nucleotides at a time. In short, mutation acts to increase—while NAGC acts to decrease—sequence divergence between paralogs. When the two sites under consideration are close-by (with respect to the NAGC mean tract length), NAGC events affecting one site are likely to incorporate the other (**Fig. 3A**). Our model makes no prior assumptions on the frequency of NAGC: unlike the tract-detection method, multiple hits are accounted for in the likelihood of the two-site model.

**Figure 3:**
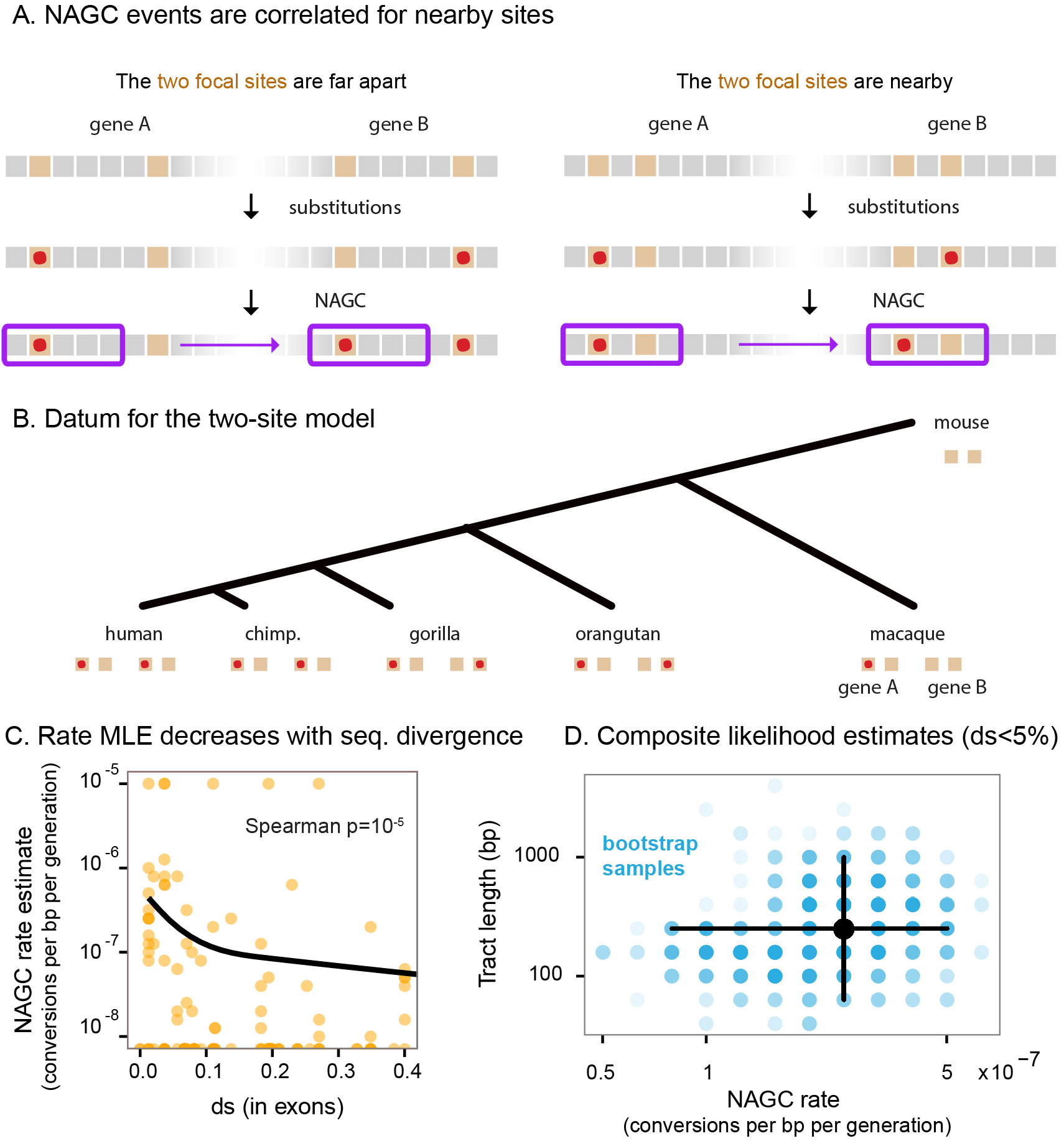
Estimation of NAGC parameters. **(A)** The two-site sequence evolution model exploits the correlated effect of NAGC on nearby sites (near with respect to the mean tract length). In this illustration, orange squares represent focal sites. Point substitutions are shown by the red points, and a converted tract is shown by the purple rectangle. **(B)** Illustration of a single datum on which we compute the full likelihood, composed of two sites in two duplicates across multiple species (except for the mouse outgroup for which only one ortholog exists). **(C)** Maximum composite likelihood (MLE) rate estimates for each intron (orange points). MLEs of zero are plotted at the bottom. The solid line shows a natural cubic spline fit. The rate decreases with sequence divergence (*ds*). We therefore only use lowly-diverged genes (*ds* ≤ 5%) to get point estimates of the baseline rate. **(D)** Composite likelihood estimates. The black point is centered at our point estimates for ds ≤ 5% genes. The blue points show 1000 non-parametric bootstrap estimates, where the intensity of each point corresponds to the number of bootstrap samplese. The corresponding 95% marginal confidence intervals are shown by black lines.

For each pair of sites in each intron in our data, we computed the likelihood of the observed alleles in all available species, over a grid of NAGC rate and mean tract length values (**Fig. 3B**). We then obtained maximum *composite* likelihood estimates (MLE) over all pairs of sites (ignoring the dependence between pairs).

We first estimated MLEs for each intron separately, and matched these estimates with ds [38] in exons of the respective gene. We found that NAGC rate estimates decrease as ds increases (Spearman *p* =1 × 10^−5^, **Fig. 3C**). This trend is likely due to a slowdown in NAGC rate, or its complete stop, as the duplicates diverge in sequence. Since our model assumes a constant NAGC rate, we concluded that the model would be most applicable to lowly diverged genes and therefore limited our parameter estimation to introns with *ds* < 5%.

We define NAGC rate as the probability that a random nucleotide is converted per basepair per generation. We estimate this rate to be 2.5 × 10^−7^ ([0.8 × 10^−7^, 5.0 × 10^−7^] 95% nonparametric bootstrap CI, **Fig. 3D**). This estimate accords with previous estimates based on smaller sample sizes using polymorphism data [26, 43] and is an order of magnitude slower than the AGC rate [70, 19]. We simultaneously estimated a mean NAGC tract length of 250bp ([63, 1000] nonparametric bootstrap CI)—consistent with estimates for AGC [30, 70]) and with a meta-analysis of many NAGC mutation accumulation experiments and NAGC-driven diseases [43].

### Live fast, stay young? The effect of NAGC on neutral sequence divergence

We next consider the implications of our results on the divergence dynamics of orthologs post duplication. In light of the high rate we infer, the question arises: if the divergence of paralogous sequences through point mutation is much slower than the elimination of divergence by NAGC [34, 56], should we expect gene duplicates never to diverge in sequence?

We considered several models of sequence divergence (**SI Appendix**). First, we considered a model where NAGC acts at the constant rate that we estimated throughout the duplicates’ evolution (“continuous NAGC”). In this case, divergence is expected to plateau around 4.5%, and concerted evolution continues for a long time (red line in **Fig. 4**; in practice there will eventually be an “escape” through a chance rapid accumulation of multiple mutations [63, 16]). However, NAGC is hypothesized to be contingent on high sequence similarity between paralogs.

**Figure 4:**
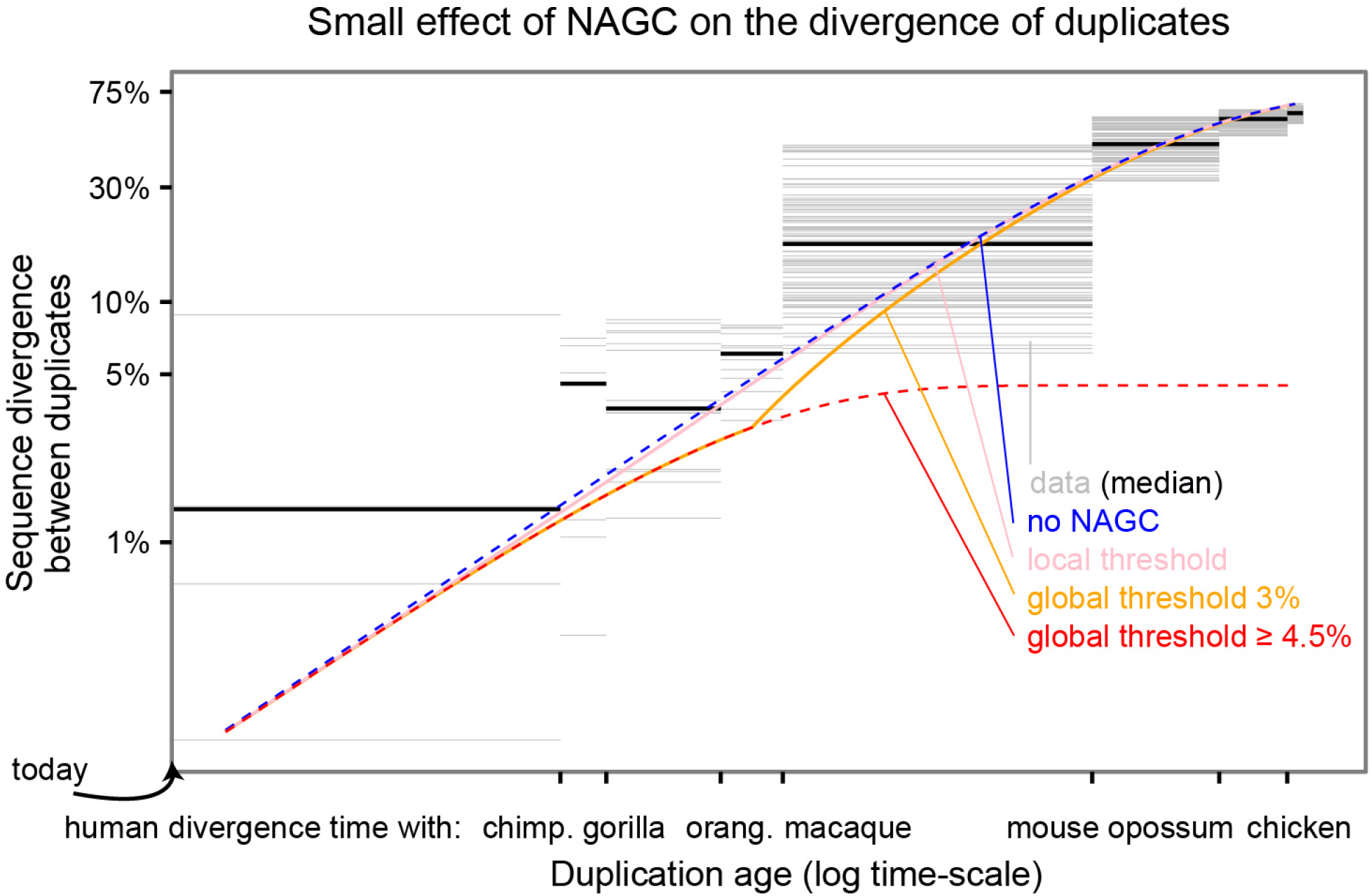
The effect of NAGC on the divergence of duplicates. The figure shows both data from human paralogs and theoretical predictions of different NAGC models. The blue line shows expected divergence in the absence of NAGC, and the red line shows the expectation with NAGC acting continuously. The pink, orange and red lines show the mean sequence divergence for models in which NAGC initation is contingent on sequence similarity between the paralogs. The grey horizontal bars correspond to human duplicate pairs. The duplication time for each pair is inferred by examining the non-human species that carry orthologs for both of the human paralogs. Y-axis shows the sequence dissimilarity between the two human paralogs.

We therefore considered two alternative models of NAGC dynamics. First, a model in which NAGC acts only while the sequence divergence between the paralogs is below some threshold (“global threshold”). Second, a model in which the initiation of NAGC at a site is contingent on perfect sequence homology at a short 400bp flanking region upstream to the site (“local threshold”, [32, 43, 11]). The local threshold model yielded a similar average trajectory to that in the absence of NAGC. A global threshold of as low as 4.5% may lead to an extended period of concerted evolution as in the continuous NAGC model. A global threshold of < 4.5% results in a different trajectory. For example, with a global threshold of 3%, duplicates born at the time of the primates most recent common ancestor (MRCA) would diverge at 3.9% of their sequence, as compared to 5.7% in the absence of NAGC (**Fig. 4; Figs. S10,S11,S12** show trajectories for other rates and threshold values).

Lastly, we asked what these results mean for the validity of molecular clocks for gene duplicates. We examined the explanatory power of different theoretical models for synonymous divergence in human duplicates. We wished to obtain an estimate of the age of duplication that is independent of *ds* between the human duplicates; we therefore used the extent of sharing of both paralogs in different species as a measure of the duplication time. For example, if a duplicate pair was found in human, gorilla and orangutan—but only one ortholog was found in macaque—we estimated that the duplication occurred at the time interval between the human-macaque split and the human-orangutan split. Except for the continuous NAGC model, all models displayed similar broad agreement with the data (**Fig. 4**).

The small effect of NAGC on divergence levels is intuitive in retrospect: for identical sequences, NAGC has no effect. Once differences start to accumulate, there is only a small window of opportunity for NAGC to act before the paralogous sequences escape from its hold. This suggests that neutral sequence divergence (e.g. ds) may be an appropriate molecular clock even in the presence of NAGC (as also suggested by [14, 13, 36]).

## Discussion

In this work, we identify recently converted regions in humans and other primates, and estimate the parameters that govern NAGC. Previously, it has been somewhat ambiguous whether concerted evolution observations were due to natural selection, pervasive NAGC, or a combination of the two [60, 41, 27]. Today, equipped with genomic data, we can revisit the pervasiveness of concerted evolution; the data in **Fig. 4** suggests that in humans, duplicates’ divergence levels are roughly as expected from the accumulation of point mutations alone. When we plugged in our estimates for NAGC rate, most mechanistic models of NAGC also predicted a small effect on neutral sequence divergence. This result suggests that neutral sequence divergence may be an appropriate molecular clock even in the presence of NAGC.

One important topic left for future investigation is the variation of NAGC parameters. Our model assumes constant action of NAGC through time and across the genome in order to get a robust estimate of the mean parameters. However, substantial variation likely exists across gene pairs due to factors such as recombination rate, sequence context, physical distance between paralogs (**Fig. S9; SI appendix**) and sequence similarity. These factors can also have very different distributions in pervasive, highly homologous sequences other than segmental gene duplicates. For example, long terminal repeats comprise several percent of the genome, and experience pervasive NAGC [66].

Our estimates for the parameters that govern the mutational mechanism alone could guide future studies of other forces guiding the evolution of gene duplicates, such as natural selection. Together with contemporary efforts to measure the effects of genomic factors on gene conversion, our results may clarify the potential of NAGC to drive disease, improve the dating of molecular events and further our understanding of the evolution of gene duplicates.

## Methods

### Gene families data

To investigate NAGC in duplicate sequences, we used a a set of 1,444 reciprocal best-matched protein-coding gene pairs in the human reference genome that we had assembled previously [36] using the human reference genome (build 37). We focused on young pairs consistent with a duplication after the human-mouse split, and identified their orthologs in the reference genomes of chimpanzee, gorilla, orangutan, macaque and mouse (**Table S1**). We focused our analysis on intronic sequences at least 50bp away from intron-exon junctions. For each of the two inference tasks we applied additional method-specific filters (**SI Appendix**)–leaving us with 75 gene families for parameter estimation and 39 gene families for inference of converted tracts.

### Two-site model

#### Transition matrix

We consider the evolution of two biallelic sites in two duplicate genes as a discrete homogeneous Markov Process. We describe these four sites with a 4-bit vector (“state vector”). The state *l_A_l_B_r_A_r_B_* ∈ {0,1}^4^ corresponds to allele *l_A_* at the “left” site in copy A, allele *l_B_* at the “left” site in copy B, allele *r_A_* at the “right” site in copy A and allele *r_B_* at the “right” site in copy B. The labels 0 and 1 are defined with respect to each site separately—the state 0000 does not mean that the left and right sites necessarily have the same allele. We first derive the (per generation) transition probability matrix. There are two possible events that may result in a transition: point mutations which occur at a rate of *µ* = 1.2 × 10^−8^ per generation [34] and NAGC. The probability of a site being converted per generation is c. We consider these mutational events to be rare and ignore terms of the order *O*(*µ^2^*), *O*(*c*^2^) and *O*(µ*c*). For example, consider the per-generation transition probability from 0110 to 0100, for two sites that are *d* bp apart. This transition can happen either through point mutation at the right site of copy A, or by NAGC from copy B to copy A involving the right site but not the left. The transition probability is therefore

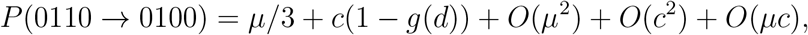

where *g(d)* is the probability of a conversion event including one of the sites given that it includes the other. Similarly, we can derive the full transition probability matrix **P**:

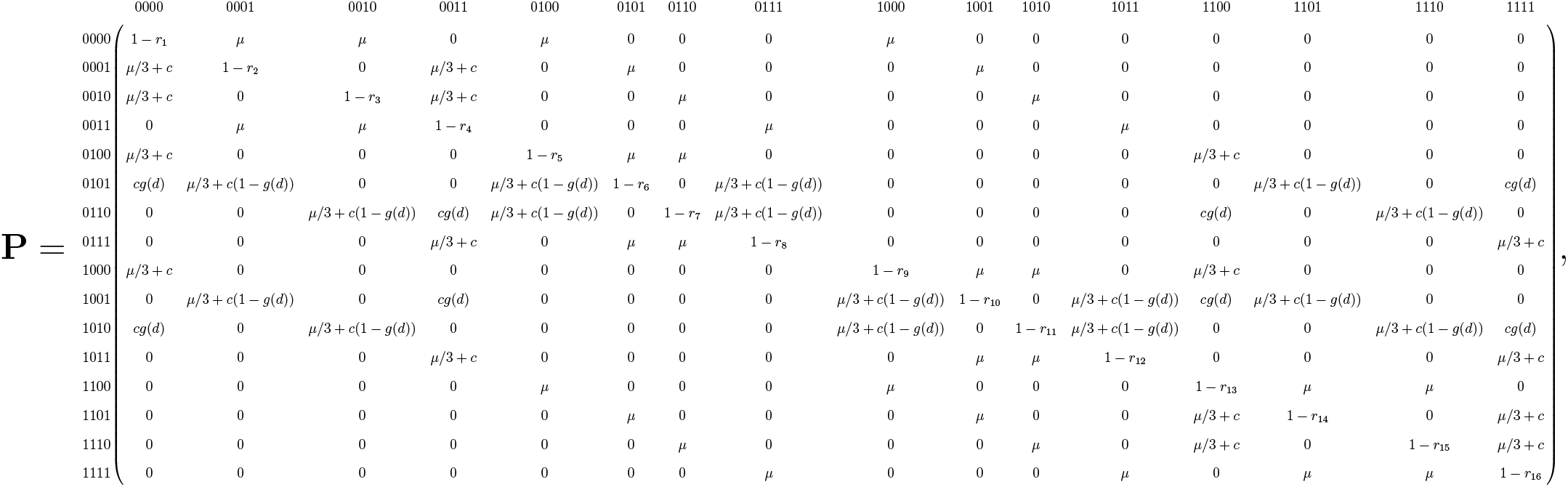

where

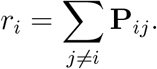

We note that this parameterization ignores possible mutations to (third and fourth) unobserved alleles.

We next derive *g(d)*. Following previous work [42], we model the tract length as geometrically distributed with mean λ. It follows that the probability of a conversion including one site conditional on it includes the other is

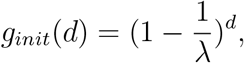

by the memorylessness of the geometric distribution. While elsewhere we assume that mutations (both point mutations and NAGC at a single site) fix at a rate equal to the mutation rate, we pause to examine this assumption for the case of a NAGC mutation including both focal sites—because the two derived alleles might decouple before one of them fixes. The probability of fixation in both sites conditional on fixation in one of them is

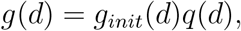

where *q*(*d*) is the probability that the second derived allele remains linked during the fixation at the first site. We make a few simplifying assumptions in evaluating *q*(*d*): The fixation time is assumed to be 4*N*_*e*_ generations where *N*_*e*_ is the (constant) effective population size. If at least one recombination event occurs, we approximate the probability of decoupling by the mean allele frequency of the first allele during fixation, 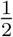. Denoting the per bp per generation recombination rate by *r*, we get:

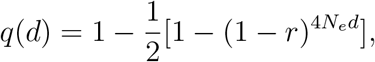

and

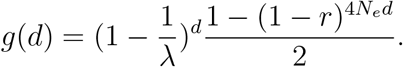

Plugging in *r* =10^−8^ [35] and *N*_*e*_ = 10^4^, we found that the probability of decoupling is high only for distances *d* where *g_init_* is already very small. Consequently, difference between *g_init_* and *g* are small throughout (**Fig. S4**). We therefore use the approximation

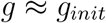

in our implementation of this model.

Lastly, we turn to compute transition probabilities along evolutionary timescales. Each datum consists of state vectors (corresponding to two biallelic sites in two paralogs) encoding the alleles in the human reference genome and 1-4 other primate reference genomes. The mouse 2-bit state (two sites in one gene) will only be used to set a prior on the root of the tree (see separate section below). We assume a constant tree—namely, a constant topology and constant edge lengths {*t_ij_*} as defined in (**Fig. S2**). We used estimates for primate split times from [54], and assumed a constant generation time of 25 years. Each node corresponds to a state. We assume that—for both mutation types—substitution occurs at a rate equal to the corresponding mutation rate. Therefore, the transition probability matrix 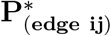 for the edge between node i and node j is

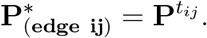

### Estimation in the two-site model

Our model describes the evolution of two sites in paralogs along primate evolution. Each of the nodes in the primate tree (**Fig. 3B**) consists of observed states—corresponding to primate references that include all four orthologous nucleotides—and hidden nodes corresponding to the state in most recent common ancestors (MRCAs) of these species. To fully determine the likelihood we must also set a prior on the state in the MRCA of all species with an observed state (“data root”). We explain the choice of prior in the **SI Appendix**.

We compute the full log likelihood for each datum (a set of 4-bit states for 2-5 primates) with transition probability matrices 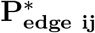. To do so in a computationally efficient way, we apply Felsenstein’s pruning algorithm [17]. We then compute the composite likelihood by summing log likelihoods over all of the data (all pairs of sites in each of the introns). We then evaluate composite likelihoods over a grid of values—the cross product of mean tract lengths 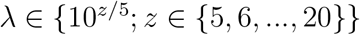 and rates 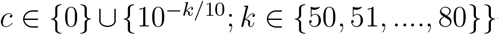—and identify the parameter values that maximize the composite likelihood.

## Acknowledgements

This work was funded by NIH grants HG008140 and MH101825 and by the Howard Hughes Medical Institute (HHMI). AH and ZG were supported in part by fellowships from the Stanford Center for Computational, Evolutionary and Human Genomics (CEHG). We thank Eilon Sharon, Doc Edge, Kelley Harris, David Knowles and Noah Rosenberg for helpful discussions.

## References

[1] Arbeithuber, B., Betancourt, A. J., Ebner, T., and Tiemann-Boege, I. Crossovers are associated with mutation and biased gene conversion at recombination hotspots. Proceedings of the National Academy of Sciences 112, 7 (2015), 2109–2114.

[2] Arguello, J. R., Chen, Y., Yang, S., Wang, W., and Long, M. Origination of an x-linked testes chimeric gene by illegitimate recombination in drosophila. PLoS Genetics 2, 5 (2006), e77.

[3] Assis, R., and Kondrashov, A. S. Nonallelic gene conversion is not GC-biased in drosophila or primates. Molecular Biology and Evolution (2011), 304.

[4] Balding, D. J., Nichols, R. A., and Hunt, D. M. Detecting gene conversion: primate visual pigment genes. Proceedings of the Royal Society of London B: Biological Sciences 249, 1326 (1992), 275–280.

[5] Betrán, E., Rozas, J., Navarro, A., and Barbadilla, A. The estimation of the number and the length distribution of gene conversion tracts from population dna sequence data. Genetics 146, 1 (1997), 89–99.

[6] Bischoff, J., Chiang, A., Scheetz, T., Stone, E., Casavant, T., Sheffield, V., and Braun, T. Genome-wide identification of pseudogenes capable of disease-causing gene conversion. Human Mutation 27, 6 (2006), 545–552.

[7] Bosch, E., Hurles, M. E., Navarro, A., and Jobling, M. A. Dynamics of a human interparalog gene conversion hotspot. Genome Research 14, 5 (2004), 835–844.

[8] Brown, D. D., and Sugimoto, K. 5s dnas of xenopus laevis and xenopus mulleri: Evolution of a gene family. Journal of Molecular Biology 78, 3 (1973), 397–415.

[9] Casola, C., Conant, G. C., and Hahn, M. W. Very low rate of gene conversion in the yeast genome. Molecular biology and evolution 29, 12 (2012), 3817–3826.

[10] Casola, C., Zekonyte, U., Phillips, A. D., Cooper, D. N., and Hahn, M. W. Interlocus gene conversion events introduce deleterious mutations into at least 1% of human genes associated with inherited disease. Genome Research 22, 3 (2012), 429–435.

[11] Chen, J.-M., Cooper, D. N., Chuzhanova, N., Férec, C., and Patrinos, G. P. Gene conversion: mechanisms, evolution and human disease. Nature Reviews Genetics 8, 10 (2007), 762–775.

[12] Dennis, M. Y., Harshman, L., Nelson, B. J., Penn, O., Cantsilieris, S., Huddleston, J., Antonacci, F., Penewit, K., Denman, L., Raja, A., et al. The evolution and population diversity of human-specific segmental duplications. Nature Ecology & Evolution 1 (2016), 0069.

[13] Dumont B. L. Interlocus gene conversion explains at least 2.7 % of single nucleotide variants in human segmental duplications. BMC Genomics (2015), 1–11.

[14] Dumont, B. L., and Eichler, E. E. Signals of historical interlocus gene conversion in human segmental duplications. PLoS One 8, 10 (2013), e75949.

[15] Duret, L., and Galtier, N. Biased gene conversion and the evolution of mammalian genomic landscapes. Annual Review of Genomics and Human Genetics 10 (2009), 285–311.

[16] Fawcett, J. A., and Innan, H. Neutral and non-neutral evolution of duplicated genes with gene conversion. Genes 2, 1 (2011), 191–209.

[17] Felsenstein, J., and Nov, N. Evolutionary trees from gene frequencies and quantitative characters : finding maximum likelihood estimates. Evolution 35, 6 (1981), 1229–1242.

[18] Galtier N. Gene conversion drives GC content evolution in mammalian histones. Trends in Genetics 19, 2 (2003), 65–68.

[19] Halldorsson, B. V., Hardarson, M. T., Kehr, B., Styrkarsdottir, U., Gylfason, A., Thorleifsson, G., Zink, F., Jonasdottir, A., Jonasdottir, A., Sulem, P., Masson, G., Thorsteinsdottir, U., Helgason, A., Kong, A., Gudbjartsson, D. F., and Stefansson, K. The rate of meiotic gene conversion varies by sex and age. Nature Genetics (2016).

[20] Hanikenne, M., Kroymann, J., Trampczynska, A., Bernal, M., Motte, P., Clemens, S., and Krämer, U. Hard selective sweep and ectopic gene conversion in a gene cluster affording environmental adaptation. PLoS Genetics 9, 8 (2013), e1003707.

[21] Hartasánchez, D. a., Vallès-Codina, O., Brasó-Vives, M., and Navarro, A. Interplay of Interlocus Gene Conversion and Crossover in Segmental Duplications Under a Neutral Scenario. G3 (Bethesda, Md.) 4, August (2014), 1479–1489.

[22] Heinen, S., Sanchez-Corral, P., Jackson, M. S., Strain, L., Goodship, J. A., Kemp, E. J., Skerka, C., Jokiranta, T. S., Meyers, K., Wagner, E., et al. De novo gene conversion in the RCA gene cluster (1q32) causes mutations in complement factor h associated with atypical hemolytic uremic syndrome. Human mutation 27, 3 (2006), 292–293.

[23] Hudson R. R. Two-locus sampling distributions and their application. Genetics 159, 4 (2001), 1805–1817.

[24] Huelsenbeck, J. P., Ronquist, F., et al. Mrbayes: Bayesian inference of phylogenetic trees. Bioinformatics 17, 8 (2001), 754–755.

[25] Hurles M. E. Gene conversion homogenizes the cmt1a paralogous repeats. BMC Genomics 2, 1 (2001), 11.

[26] Innan H. A two-locus gene conversion model with selection and its application to the human rhce and rhd genes. Proceedings of the National Academy of Sciences 100, 15 (2003), 8793–8798.

[27] Innan, H., and Kondrashov, F. The evolution of gene duplications: classifying and distinguishing between models. Nature Reviews Genetics 11, 2 (2010), 97–108.

[28] Jackson, M. S., Oliver, K., Loveland, J., Humphray, S., Dunham, I., Rocchi, M., Viggiano, L., Park, J. P., Hurles, M. E., and Santibanez-Koref, M. Evidence for widespread reticulate evolution within human duplicons. The American Journal of Human Genetics 77, 5 (2005), 824–840.

[29] Jakobsen, I. B., Wilson, S. R., and Easteal, S. The partition matrix: exploring variable phylogenetic signals along nucleotide sequence alignments. Molecular biology and evolution 14, 5 (1997), 474–484.

[30] Jeffreys, A. J., and May, C. A. Intense and highly localized gene conversion activity in human meiotic crossover hot spots. Nature Genetics 36, 2 (2004), 151–156.

[31] Ji, X., Griffing, A., and Thorne, J. L. A phylogenetic approach finds abundant interlocus gene conversion in yeast. Molecular Biology and Evolution 33, 9 (2016), 2469–2476.

[32] Jinks-Robertson, S., Michelitch, M., and Ramcharan, S. Substrate length requirements for efficient mitotic recombination in saccharomyces cerevisiae. Molecular and Cellular Biology 13, 7 (1993), 3937–3950.

[33] Jinks-Robertson, S., and Petes, T. D. High-frequency meiotic gene conversion between repeated genes on nonhomologous chromosomes in yeast. Proceedings of the National Academy of Sciences 82, 10 (1985), 3350–3354.

[34] Kong, A., Frigge, M. L., Masson, G., Besenbacher, S., Sulem, p., Magnusson, G., Gudjonsson, S. A., Sigurdsson, A., Jonasdottir, A., Jonasdottir, A., et al. Rate of de novo mutations and the importance of father/’s age to disease risk. Nature 488, 7412 (2012), 471–475.

[35] Kong, A., Thorleifsson, G., Gudbjartsson, D. F., Masson, G., Sigurdsson, A., Jonasdottir, A., Walters, G. B., Jonasdottir, A., Gylfason, A., Kristinsson, K. T., et al. Fine-scale recombination rate differences between sexes, populations and individuals. Nature 467, 7319 (2010), 1099–1103.

[36] Lan, X., and Pritchard, J. K. Coregulation of tandem duplicate genes slows evolution of subfunctionalization in mammals. Science 352, 6288 (2016), 1009–1013.

[37] Lesecque, Y., Mouchiroud, D., and Duret, L. GC-biased gene conversion in yeast is specifically associated with crossovers: molecular mechanisms and evolutionary significance. Molecular biology and evolution 30, 6 (2013), 1409–1419.

[38] Li, W.-H., and Graur, D. Fundamentals of molecular evolution. Sinauer Associates, 1991.

[39] Lichten, M., and Haber, J. Position effects in ectopic and allelic mitotic recombination in saccharomyces cerevisiae. Genetics 123, 2 (1989), 261–268.

[40] Lorson, C. L., Hahnen, E., Androphy, E. J., and Wirth, B. A single nucleotide in the smn gene regulates splicing and is responsible for spinal muscular atrophy. Proceedings of the National Academy of Sciences 96, 11 (1999), 6307–6311.

[41] Mano, S., and Innan, H. The evolutionary rate of duplicated genes under concerted evolution. Genetics 180, 1 (2008), 493–505.

[42] Mansai, S. P., and Innan, H. The power of the methods for detecting interlocus gene conversion. Genetics 184, 2 (2010), 517–527.

[43] Mansai, S. P., Kado, T., and Innan, H. The rate and tract length of gene conversion between duplicated genes. Genes 2, 2 (2011), 313–331.

[44] McGrath, C. L., Casola, C., and Hahn, M. W. Minimal effect of ectopic gene conversion among recent duplicates in four mammalian genomes. Genetics 182, 2 (2009), 615–622.

[45] McVean, G., Awadalla, P., and Fearnhead, P. A coalescent-based method for detecting and estimating recombination from gene sequences. Genetics 160, 3 (2002), 1231–1241.

[46] Mitchell M. B. Aberrant recombination of pyridoxine mutants of neurospora. Proceedings of the National Academy of Sciences 41, 4 (1955), 215–220.

[47] Narasimhan, V. M., Rahbari, R., Scally, A., Wuster, A., Mason, D., Xue, Y., Wright, J., Trembath, R. C., Maher, E. R., van Heel, D. A., et al. Estimating the human mutation rate from autozygous segments reveals population differences in human mutational processes. Nature Communications 8 (2017).

[48] Nei M. Molecular evolutionary genetics. Columbia university press, 1987.

[49] Odenthal-Hesse, L., Berg, I. L., Veselis, A., Jeffreys, A. J., and May, C. A. Transmission distortion affecting human noncrossover but not crossover recombination: a hidden source of meiotic drive. PLoS Genetics 10, 2 (2014), e1004106.

[50] Ohta T. How gene families evolve. Theoretical Population Biology 37, 1 (1990), 213–219.

[51] Roesler, J., Curnutte, J. T., Rae, J., Barrett, D., Patino, P., Chanock, S. J., and Goerlach, A. Recombination events between the p47-phoxgene and its highly homologous pseudogenes are the main cause of autosomal recessive chronic granulomatous disease. Blood 95, 6 (2000), 2150–2156.

[52] Rozen, S., Skaletsky, H., Marszalek, J. D., Minx, P. J., Cordum, H. S., Waterston, R. H., Wilson, R. K., and Page, D. C. Abundant gene conversion between arms of palindromes in human and ape Y chromosomes. Nature 423, 6942 (2003), 873–876.

[53] Sawyer S. Statistical tests for detecting gene conversion. Molecular Biology and Evolution 6, 5 (1989), 526–538.

[54] Scally, A., Dutheil, J. Y., Hillier, L. W., Jordan, G. E., Goodhead, I., Herrero, J., Hobolth, A., Lappalainen, T., Mailund, T., Marques-Bonet, T., McCarthy, S., Montgomery, S. H., Schwalie, P. C., Tang, Y. A., Ward, M. C., Xue, Y., Yngvadottir, B., Alkan, C., Andersen, L. N., Ayub, Q., Ball, E. V., Beal, K., Bradley, B. J., Chen, Y., Clee, C. M., Fitzgerald, S., Graves, T. A., Gu, Y., Heath, P., Heger, A., Karakoc, E., Kolb-Kokocinski, A., Laird, G. K., Lunter, G., Meader, S., Mort, M., Mullikin, J. C., Munch, K., O’Connor, T. D., Phillips, A. D., Prado-Martinez, J., Rogers, A. S., Sajjadian, S., Schmidt, D., Shaw, K., Simpson, J. T., Stenson, P. D., Turner, D. J., Vigilant, L., Vilella, A. J., Whitener, W., Zhu, B., Cooper, D. N., de Jong, P., Dermitzakis, E. T., Eichler, E. E., Flicek, P., Goldman, N., Mundy, N. I., Ning, Z., Odom, D. T., Ponting, C. P., Quail, M. A., Ryder, O. A., Searle, S. M., Warren, W. C., Wilson, R. K., Schierup, M. H., Rogers, J., Tyler-Smith, C., and Durbin, R. Insights into hominid evolution from the gorilla genome sequence. Nature 483, 7388 (2012), 169–75.

[55] Schildkraut, E., Miller, C. A., and Nickoloff, J. A. Gene conversion and deletion frequencies during double-strand break repair in human cells are controlled by the distance between direct repeats. Nucleic Acids Research 33, 5 (2005), 1574–1580.

[56] Ségurel, L., Wyman, M. J., and Przeworski, M. Determinants of mutation rate variation in the human germline. Annual Review of Genomics and Human Genetics 15 (2014), 47–70.

[57] Smith G. P. Unequal crossover and the evolution of multigene families. In Cold Spring Harbor Symposia on Quantitative Biology (1974), vol. 38, Cold Spring Harbor Laboratory Press, pp. 507–513.

[58] Smith, G. P., Hood, L., and Fitch, W. M. Antibody diversity. Annual Review of Biochemistry 40, 1 (1971), 969–1012.

[59] Storz, J. F., Baze, M., Waite, J. L., Hoffmann, F. G., Opazo, J. C., and Hayes, J. P. Complex signatures of selection and gene conversion in the duplicated globin genes of house mice. Genetics 177, 1 (2007), 481–500.

[60] Sugino, R. P., and Innan, H. Selection for more of the same product as a force to enhance concerted evolution of duplicated genes. Trends in Genetics 22, 12 (2006), 642–644.

[61] Taghian, D. G., and Nickoloff, J. A. Chromosomal double-strand breaks induce gene conversion at high frequency in mammalian cells. Molecular and Cellular Biology 17, 11 (1997), 6386–6393.

[62] Takuno, S., and Innan, H. Selection to maintain paralogous amino acid differences under the pressure of gene conversion in the heat-shock protein genes in yeast. Molecular Biology and Evolution 26, 12 (2009), 2655–2659.

[63] Teshima, K. M., and Innan, H. The effect of gene conversion on the divergence between duplicated genes. Genetics 166, 3 (2004), 1553–1560.

[64] Teshima, K. M., and Innan, H. Neofunctionalization of duplicated genes under the pressure of gene conversion. Genetics 1398, March (2007), 1385–1398.

[65] Thornton, K., and Long, M. Excess of amino acid substitutions relative to polymorphism between x-linked duplications in drosophila melanogaster. Molecular Biology and Evolution 22, 2 (2005), 273–284.

[66] Trombetta, B., Fantini, G., D’Atanasio, E., Sellitto, D., and Cruciani, F. Evidence of extensive non-allelic gene conversion among ltr elements in the human genome. Scientific reports 6 (2016), 28710.

[67] Watnick, T. J., Gandolph, M. A., Weber, H., Neumann, H. P., and Germino, G. G. Gene conversion is a likely cause of mutation in pkd1. Human Molecular Genetics 7, 8 (1998), 1239–1243.

[68] Weiller G. F. Phylogenetic profiles: a graphical method for detecting genetic recombinations in homologous sequences. Molecular Biology and Evolution 15, 3 (1998), 326–335.

[69] Whelden Cho, J., Khalsa, G. J., and Nickoloff, J. A. Gene-conversion tract directionality is influenced by the chromosome environment. Current Genetics 34, 4 (1998), 269–279.

[70] Williams, A. L., Genovese, G., Dyer, T., Altemose, N., Truax, K., Jun, G., Patterson, N., Myers, S. R., Curran, J. E., Duggirala, R., Blangero, J., Reich, D., and Przeworski, M. Non-crossover gene conversions show strong GC bias and unexpected clustering in humans. Elife 4 (2015), e04637.

[71] Yang, D., and Waldman, A. S. Fine-resolution analysis of products of intrachromosomal homeologous recombination in mammalian cells. Molecular and Cellular Biology 17, 7 (1997), 3614–3628.

[72] Zimmer, E., Martin, S., Beverley, S., Kan, Y., and Wilson, A. C. Rapid duplication and loss of genes coding for the alpha chains of hemoglobin. Proceedings of the National Academy of Sciences 77, 4 (1980), 2158–2162.

